# Learning cellular morphology with neural networks

**DOI:** 10.1101/364034

**Authors:** Philipp J Schubert, Sven Dorkenwald, Michał Januszewski, Viren Jain, Joergen Kornfeld

## Abstract

Reconstruction and annotation of volume electron microscopy data sets of brain tissue is challenging, but can reveal invaluable information about neuronal circuits. Significant progress has recently been made in automated neuron reconstruction, as well as automated detection of synapses. However, methods for automating the morphological analysis of nanometer-resolution reconstructions are less established, despite their diverse application possibilities. Here, we introduce cellular morphology neural networks (CMNs), based on multi-view projections sampled from automatically reconstructed cellular fragments of arbitrary size and shape. Using unsupervised training we inferred morphology embeddings (“Neuron2vec”) of neuron reconstructions and trained CMNs to identify glia cells in a supervised classification paradigm which was then used to resolve neuron reconstruction errors. Finally, we demonstrate that CMNs can be used to identify subcellular compartments and the cell types of neuron reconstructions.

## Introduction

Advances in volume electron microscopy (VEM) have led to increasingly large 3D images of brain tissue, making manual analysis infeasible^1^. Multi-beam scanning electron microscopes^2^ and transmission electron microscopes equipped with fast camera arrays can now generate data sets exceeding 100 TB^3^, a development which was fortunately accompanied by substantial progress in neuron reconstruction ^4–9^ and the automatic analysis of synapses ^10–13^. These advances enable now automatic morphology analyses on the neuron (fragment) scale, which were previously restricted to direct segmentation error detection ^5,14^, or used manual skeletons with data-specific hand-crafted features ^11,15,16^. Cell types, and other biological properties that can be inferred from the morphology of a neuron, are required for the interpretation of a connectome^11^ and can also constrain automatic neuron reconstruction itself^17^.

Outside of the field of connectomics, many approaches have been developed for automated 3D shape analysis, including multi-view 2D projection based neural network models ^18,19^ as well as voxel- and point cloud-occupancy based 3D networks ^20,21^. Interestingly, projection based models appear to often outperform 3D architectures, possibly because of higher-resolution input due to decreased model complexity^18^.

Here, we present cellular morphology neural networks (CMNs), which use multi-view projections to enable the supervised and unsupervised analysis of cell fragments of arbitrary size while retaining high-resolution. First, we demonstrate that CMNs can be used to automate morphology feature extraction itself, by inferring low-dimensional embeddings, dubbed “Neuron2vec”, through unsupervised triplet-loss training ^22,23^. Second, we apply them to the supervised classification of glia cells and use these data to demonstrate the feasibility and effectivity of a simple top-down false merger resolution strategy. Third, we identify neuronal cell types and compartments, outperforming methods with hand-crafted features based on skeleton representations on the same data, and finally perform high-resolution cell surface segmentation to identify dendritic spines.

## Results

### Cellular morphology learning networks

CMNs are convolutional neural networks (CNNs) optimized for the analysis of multi-channel 2D projections of cell reconstructions, inspired by multi-view CNNs for the classification of objects fitting into projections from one rendering site ^18,19^.

Su et al.^18^ rendered views of an entire object, potentially sacrificing crucial detail when applied to reconstructions of entire neurons (or very large objects in general), which can have processes as thin as 50 nm extending over millimeters^24^. To address this problem, we used a sampling algorithm that homogeneously probes an entire cell at many locations (Methods; Fig. 1a,b) that are then analyzed either individually or in combination by a CMN. The used neuron reconstructions were from a songbird basal ganglia data set and consisted of flood-filling network (FFN) created supervoxels^25^ (SV), which were agglomerated to sets of super-SVs (SSV)^8^, each corresponding to a single neuron. A rectangular field of view (FoV) was chosen for the projection to capture the elongated shape of most neuronal arbors better. Additionally, we incorporated image channels beyond the cell shape (here represented through depth-map projections) of the same rendering perspective. This extension allows the CMN to analyze the geometry and density of objects contained in a cell, such as mitochondria and other organelles (Fig. 1c-d).

We implemented the view rendering engine with OpenGL and the neural network models using the ElektroNN neural network library (www.elektronn.org). The models were trained using standard loss optimization procedures (Adam or stochastic gradient descent) on various supervised and unsupervised tasks, which are described in the following to demonstrate the versatility of the approach.

**Fig. 1.**
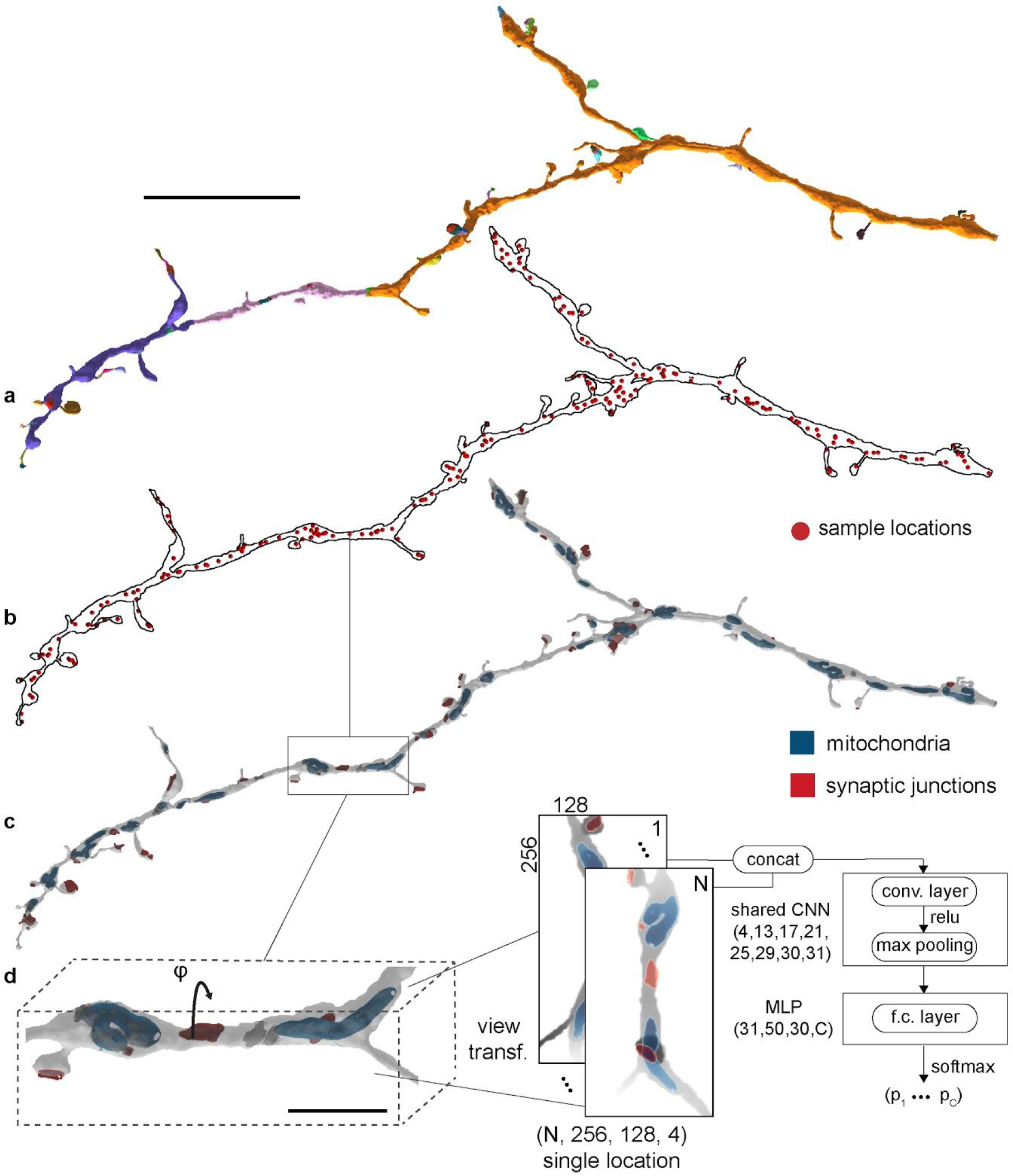
Multi-view generation for a given neurite reconstruction and model architecture. **a** FFN reconstruction and agglomeration of a dendrite, with SVs individually colored. **b** Sampled multi-view projection locations indicated as red spheres. **c** 2D projection of the whole neurite reconstruction with cell organelles (blue: mitochondria; red: synaptic junctions). **d** Each location was rendered from two different perspectives, one orthogonal to the 1st and 2^nd^ principal component (p.c.), the other one rotated by φ around the 1st p.c., i.e. orthogonal to the 1st and 3rd p.c. (here: φ = 90°). The model architecture for the general N-view case consisted of seven 2D CNN layers that extract view features, three fully connected layers (MLP), and a final softmax layer that computes class-wise probabilities (C classes). Scale bars are 10 μm in **a** and 2 μm in **d**.

### Neuron2Vec embedding

Supervised models require often hard to obtain manual ground truth, making alternative objective functions and models based on intuitive isomorphisms of the underlying data desirable. We trained a CMN using triplet loss^23^ to learn a latent space (embedding; dimension d_z_=10) of single renderings based on prior information about the similarity of three inputs *x_ref_*, *x*_+_ and *x*_−_. At every location two renderings (see above; φ = 50°) were generated which served as similar pair (*x_ref_* and *x*_+_). In contrast, a single rendering at a different, randomly sampled location was used as the dissimilar example *x*_−_ (Methods). During training, the views were sampled from 372 cell or cell fragment reconstructions (181.02 mm; 19.95 giga voxel (GV); 32,324 μm^3^).

We explored the information content of the embedding through inspection of clusters in a 2D t-SNE^26^ projection (Fig. 2a) of an example cell reconstruction (Fig. 2b) and by fitting a k-nearest neighbor classifier (kNN; k=5; uniform weights) to the Neuron2vec encoding extracted from a set of neurites with cellular compartment ground truth annotations (same as in “Cellular compartment identification”; total path length: 30.16 mm; 3.05 GV; 4947 μm^3^).

The predictive performance of the kNN classifier on the triplet-CMN (t-CMN) latent vectors of a cellular compartment validation set (2.73 mm; 2.48 GV; 4023 μm^3^) was particularly low for dendrites (F1-score for dendrite: 0.529, axon: 0.791, soma: 0.923). We were able to increase the classification performance when we allowed to draw the similar view from close-by rendering locations during training as well (two and eight additional rendering locations N_r_=3 and 9), which had a smoothing effect across the inferred latent vectors of adjacent rendering locations (Fig. S2b; F1-score for dendrite, axon, soma with N_r_=3: 0.639, 0.843, 0.951; N_r_=9: 0.831, 0.911, 0.960).

Axons, or thin processes in general, and soma regions could be readily identified based on a color-projection of the embedding using principal component analysis (PCA, Fig. 2 and S2), whereas, for example, spiny dendrites showed a heterogeneous and less continuous spectrum (inset Fig. 2b). Interestingly, views did not only group by compartment type, but also formed sub-clusters separating views with cell objects (such as mitochondria and synaptic junctions) from those without (Fig. 2c i,ii).

We next took a supervised end-to-end approach and examined ground truth based CMN models.

**Fig. 2.**
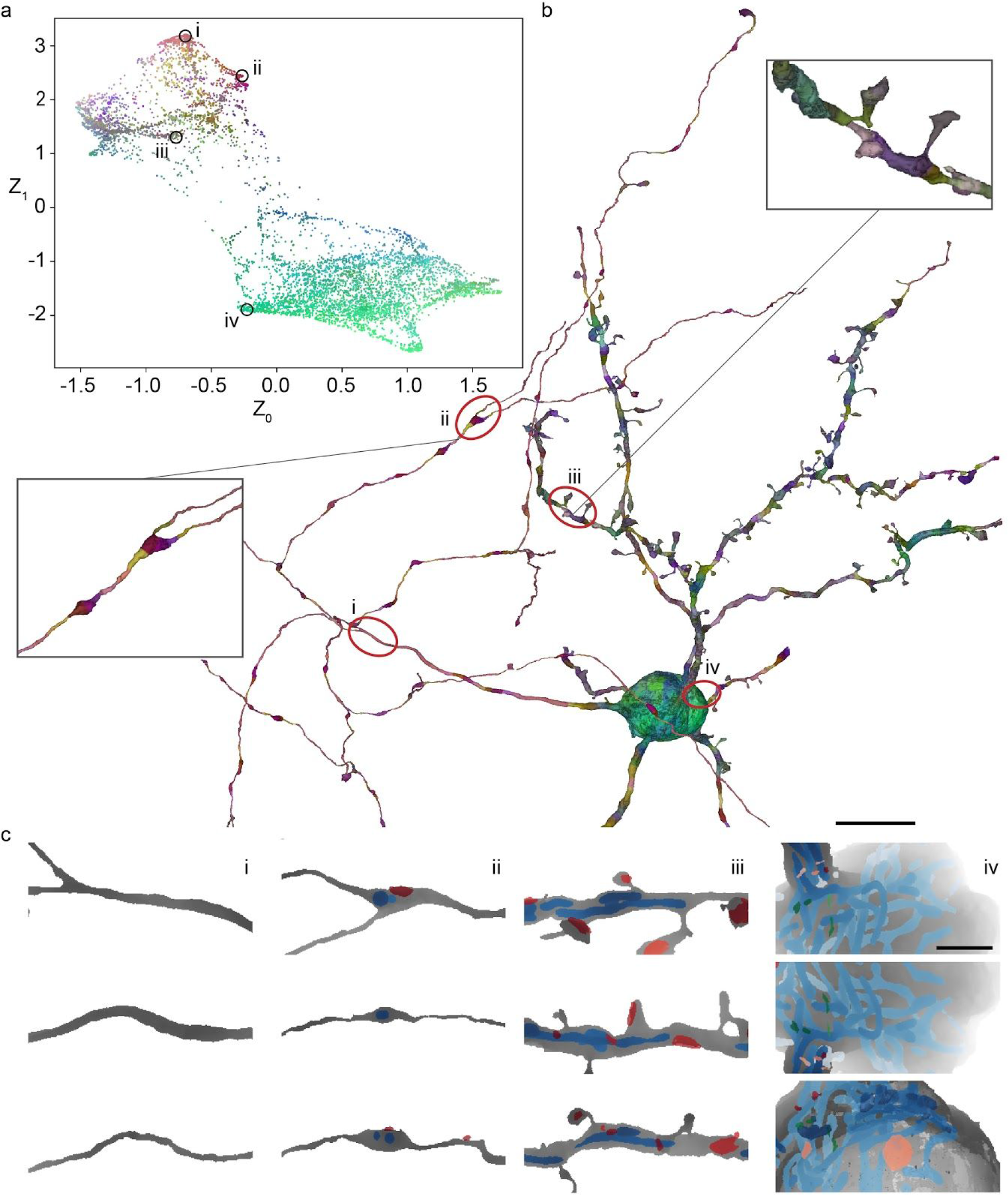
Neuron2Vec embeddings learned by a t-CMN (d_z_=10; N_r_=1; top three p.c.’s covered 60.2% of the variance and were used as RGB values). **a** t-SNE transformed latent space of an example cell reconstruction, shown in **b**(coloring as in **a**). Insets show close-ups of an axonal bouton and a spiny dendrite. **c** Example views of axonal segments (i), axonal segments with mitochondria (blue) or synaptic junctions (red) (ii), spiny dendrite segments (iii) and at the soma (iv; green: vesicle clouds). View order is determined by the distance to the plot locations shown in **a**. Scale bars in **b** and **c** are 10 μm and 2 μm.

### Glia detection and top-down segmentation error correction

Due to the dense heavy-metal staining used for VEM data, the acquired images contain all cell types, including astrocytes and other glia types, which are usually not considered for connectomic analysis. Astrocytes have tight surface contacts with neurons to supply them with nutrients and regulate the local environment^27^, making astrocyte-neurite mergers a problem in automated reconstruction pipelines. While it can be difficult for humans to determine from the raw EM data whether a process belongs to a glia cell (Fig. S1), it seemed straightforward to solve this classification task using the established multi-view representation, due to their distinct shape (Fig. 3a). A CMN was trained and validated on a set of manually annotated SVs (from 34 neurite reconstructions and 118 glia SVs; 368.74 mm; 16.48 GV; 26,695 μm^3^), achieving an average F1-score of 0.979 (precision: 0.985; recall: 0,974; N=9,695 multi-views; Fig. S3). We then explored the effect of varying data context and view resolution on classification performance and found that, as expected, context is crucial (largest context tested: 8 × 4 × 4 μm^3^; F1-score of 0.900 upon 75% reduction; Fig. S3), while reduced resolution barely affects the performance of CMNs (F1-score of 0.967 upon 75% reduction). In agreement with this, SVs (N_Neuron_: 84; N_Glia_: 85; 27 μm^3^) part of larger automatic neurite reconstructions, i.e. with a bounding box diagonal (BBD) above 8 μm, showed a significantly better performance per SV (F1 score 0.964) in comparison to all SVs (F1 score 0.868; Fig. 3b).

Can we exploit the excellent glia classification results to reduce the rate of false mergers in a segmentation? The naive solution would be the simple removal of all SV classified as glia fragments, which could however induce large-scale false splits in the case of false positive classifications (Fig. S5f). We therefore developed a more sophisticated splitting heuristic, ensuring that large neuron fragments remained connected in the presence of small misclassifications.

The agglomeration of SVs can be represented as a graph, connecting adjacent SVs (nodes) of the same SSV (neurite) with edges. In this graph glia predictions were used as node properties to identify connected glia and neuron components with their respective sizes. Sufficiently big (BBD ≥ 8.0 μm) glia components were removed and added to the glia graph, while small glia components (BBD < 8.0 μm) remained in the neuron graph. The resulting connected components of the graphs were stored as individual glia and neuron SSVs, respectively (Fig. 3d). We evaluated the SV labels of 12 neurite reconstructions with predicted glia mergers (N_Neuron_: 321 SV; N_Glia_: 295 SV; 4181 μm^3^; Fig. S5) after the splitting procedure and found a significantly better classification performance when including the SV volume compared to unit weights (weighted F1-score of 0.995 vs. 0.937; Fig. 3e). Using this approach 882 splits were introduced in 181 neurites with a total of 3,866,049 SVs classified as neuron (606,922 μm^3^) and 2,154,960 as glia (110,342 μm^3^), yielding an astrocyte volume fraction of about 0.154. It should be noted that the EM data set was prepared in a way to preserve the ECS, which may affect the morphology of the glial processes^28^.

We then attempted to reconstruct the 27 astrocytes identified in the data set starting at their somata and assigning each glial fragment to the closest astrocyte soma (see Methods), justified by reports that astrocytes establish roughly spherical, non-overlapping territories^29^. While the CMN-based glia identification appears in principle promising (see Fig. S6 for an analysis of automatically extracted glia-blood vessel contacts), our approach likely merged arbors from other glia cells, due to them reaching into the data set from the outside.

**Fig. 3.**
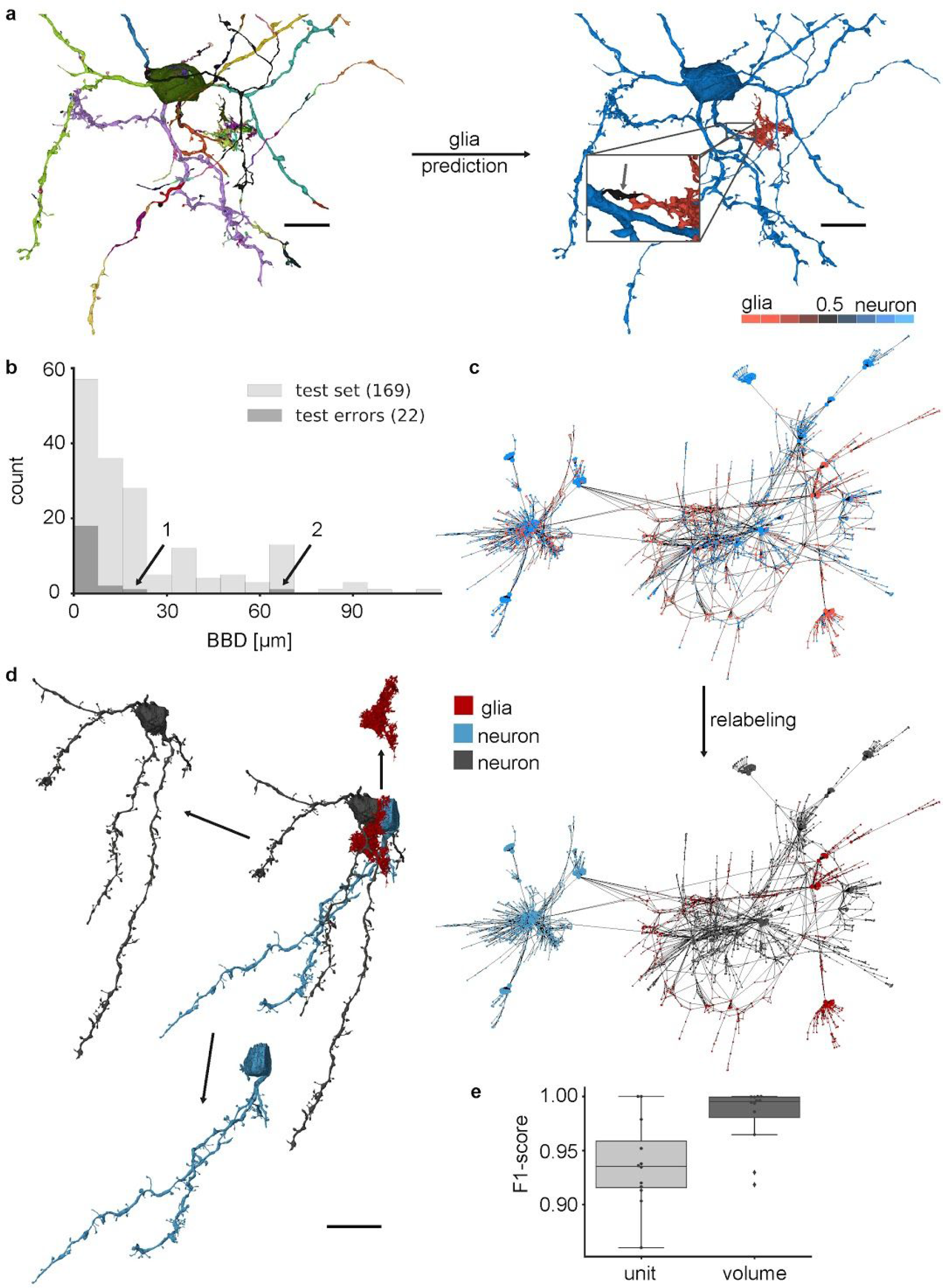
Glia prediction and reconstruction splitting procedure. **a** Each SV (unique coloring) of a neurite reconstruction was classified as glia (red) or neuron (blue) based on the CMN output. **b** Corresponding neurite length (BBDs) distribution for 169 randomly drawn SVs (light grey) vs. SVs with classification errors (dark grey). Arrows indicate errors of SVs with a BBD ≥ 10 μm (see Fig. S4 for example renderings). **c** SSV-graph representation (spring-layout) of a glia-predicted neurite reconstruction (SV volume represented by node size). **d** Connected component (CC) meshes and SSV graph indicating CCs after the splitting procedure. **e** Boxplot (box: mean, Q1-Q3; whiskers: 1.5 IQR) of SV classification performance with unit weights (light grey; mean: 0.938; sample standard deviation, s.d.: 0.039) and volume weighted (dark grey; mean: 0.981; s.d.: 0.027) splitting performance of 12 neurites (significance of Wilcoxon rank-sum test: 20 μm for **d**. *p* = 0.028 < α = 0.05 with N=12). Scale bars are 10 μm for **a**, **b** and 20 μm for **d**.

### Neuron type classification

Similar to glia cells, many neuronal types can be identified based on their morphology^15^, a feature that was used well before connectivity-based methods^30^ and other approaches (reviewed in ^31^). We recently demonstrated on the same songbird data set that a feature-based method with random forest classifiers (RFCs) can be used to identify the four main cell types, excitatory axons (EA), medium spiny neurons (MSN), pallidal-like neurons (GP) and interneurons (INT)^11^. However, manual neurite skeletons, as well as hand-designed feature vectors were required as basis for this classification method.

In contrast to the classification of astrocytes, many neuronal types are indistinguishable when using only a local view (i.e. spatially focused) of a neurite (F1-scores for N=1: 0.885 and after majority vote: 0.884; training set: 145.98 mm, 17.21 GV, 27,872 μm^3^; test set: 65.04 mm, 7.31 GV, 11,843 μm^3^). A simple solution would be to increase the FoV for a single view, capturing the entire extent of a neuron, similar to the proposed object classification method by Su et al.^18^. However, this approach reduces the resolution for a given view-size, obscuring potentially important details. Alternatively, local views from different locations can be sampled at random and combined to a global representation (multi-views of size N, further called N-views; Fig. 4a) by the neural network model. This approach does not sacrifice resolution and increased the classification performance substantially (N=10 views: F1-score of 0.987, and 0.968 on SSV; Fig. 4b). It should be noted that lower resolution, zoomed out views might still be beneficial, which was albeit not explored further by us. Interestingly, a greater number of sample locations did not necessarily increase performance (F1-score for N=60: 0.984, SSV: 0.966; Fig. 4b), possibly due to increasing model complexity and decreasing diversity of the training data through fewer independent view samples per neuron. This result was consistent with the observation that the N-view F1-scores throughout all Ns were significantly lower without shuffling (views were analyzed in the order they were created) (Fig. 4c), which led to spatially correlated N-view sets, i.e. they likely consisted of views from a single compartment type only. For all N, additional models were trained to predict sets of views without cell organelle channels, which, in agreement with our previous results using RFCs^11^, reduced the F1-score substantially (e.g. F1-score reduction of 0.10 for N=20; Fig. 4b). Interestingly, cell organelles have shown to be crucial for the correct cell type prediction of reconstructions that contained only few N-views (Fig. 4b).

**Fig. 4.**
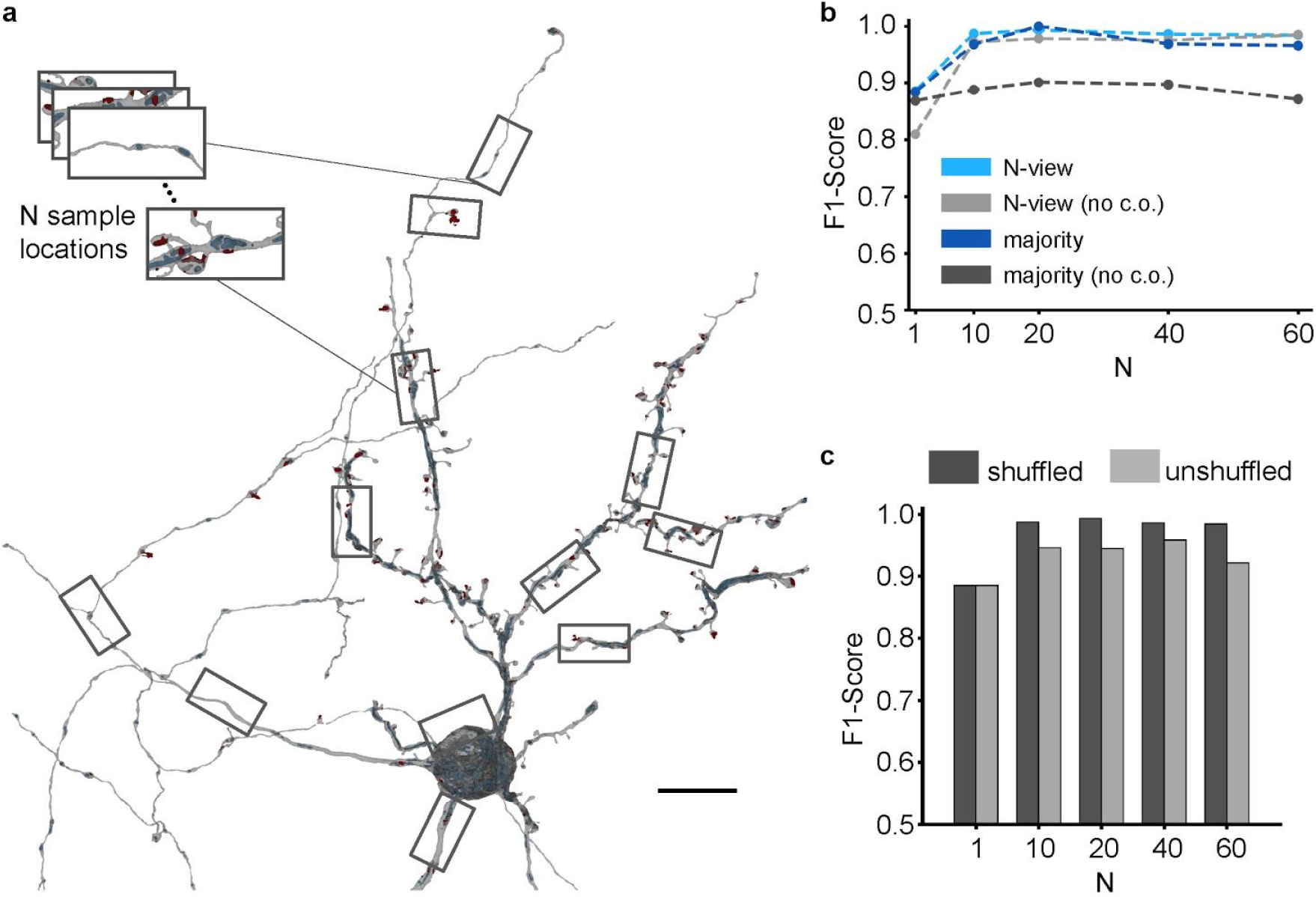
Cell type inference using CMNs. **a** N-view fingerprint of a neuron reconstruction. **b** Weighted F1-score of cell types with and without cell organelles (no c.o.). Light grey and light blue represent performance of individual N-views (class support for N=1: EA: 16,774, MSN: 90,662, GP: 3958, INT: 8806; N=10: EA: 1658, MSN: 9048, GP: 394, INT: 880; N=20: EA: 812, MSN: 4514, GP: 196, INT: 440; N=40: EA: 388, MSN: 2245, GP: 97, INT: 220; N=60: EA: 251, MSN 1489, GP: 65, INT: 146); dark tones represent the average F1-score on neurite level after majority voting (class support EA: 60, MSN: 39). **c** F1-score for shuffled (dark grey) and unshuffled (light grey) N-view stacks (class support as in **b**). Scale bar is 10 μm.

### Cellular compartment identification

We next attempted to analyze the FFN neuron segmentation by identifying subcellular compartments (axon, dendrite and soma) at single multi-view locations (Fig. 5a).

Similar to the glia model, a reduction of the original FoV of 8 × 4 × 4 μm^3^ to 2 × 1 × 1 μm^3^ reduced the performance (F1-score on validation set of 0.996 with full res. vs. 0.913), while a 4-fold downsampling of the multi-views had almost no effect (reduction of 0.014, Fig. 5b). Exclusion of the cell organelle information at full resolution reduced the F1-score by 0.085. Not surprisingly, the performance of the soma class was barely affected by this or any other changes to the input, whereas the discrimination between axons and dendrites was strongly dependent on cell organelle information (Fig. 5a).

The performance of the CMN approaches were directly compared with the skeleton-based RF classification developed before by us^11^ on a set of 28 manually annotated reconstructions (20.75 mm; 1.31 GV; 2130 μm^3^). Two different FoVs (implemented as maximum skeleton traversal distances for the RFC approach, RFC-4, with 4 μm and RFC-8 with 8 μm) were tested with morphology features extracted from these FoVs (see Methods) and fitted to the same training data as the CMN (path length 30.16 mm; 3.05 GV; 4947 μm^3^). We further evaluated the RFC models (RFC*-4, RFC*-8) on test data including entire axons and dendrites, to exclude that performance differences originate from the slightly different window-size selection methods (skeleton traversal distance vs. multi-view FoV) or fragmented reconstructions of somata (Fig. S7d). Although the CMN (2 views; Fig. 5b) was operating on less context (multi-views were based on a 8 μm × 4 μm × 4 μm subvolume vs. a maximum possible subvolume of 16 μm × 16 μm × 16 μm for the RFC-8 model), it outperformed both skeleton-based models and the Neuron2Vec-kNN classifier (t-CMN; Fig 6c). Local mis-classifications could be corrected successfully by applying a sliding window majority vote (Fig. S7a,b). Increasing the latent space dimension of the t-CMN to 25 did not significantly affect its performance (Fig. S7).

**Fig. 5.**
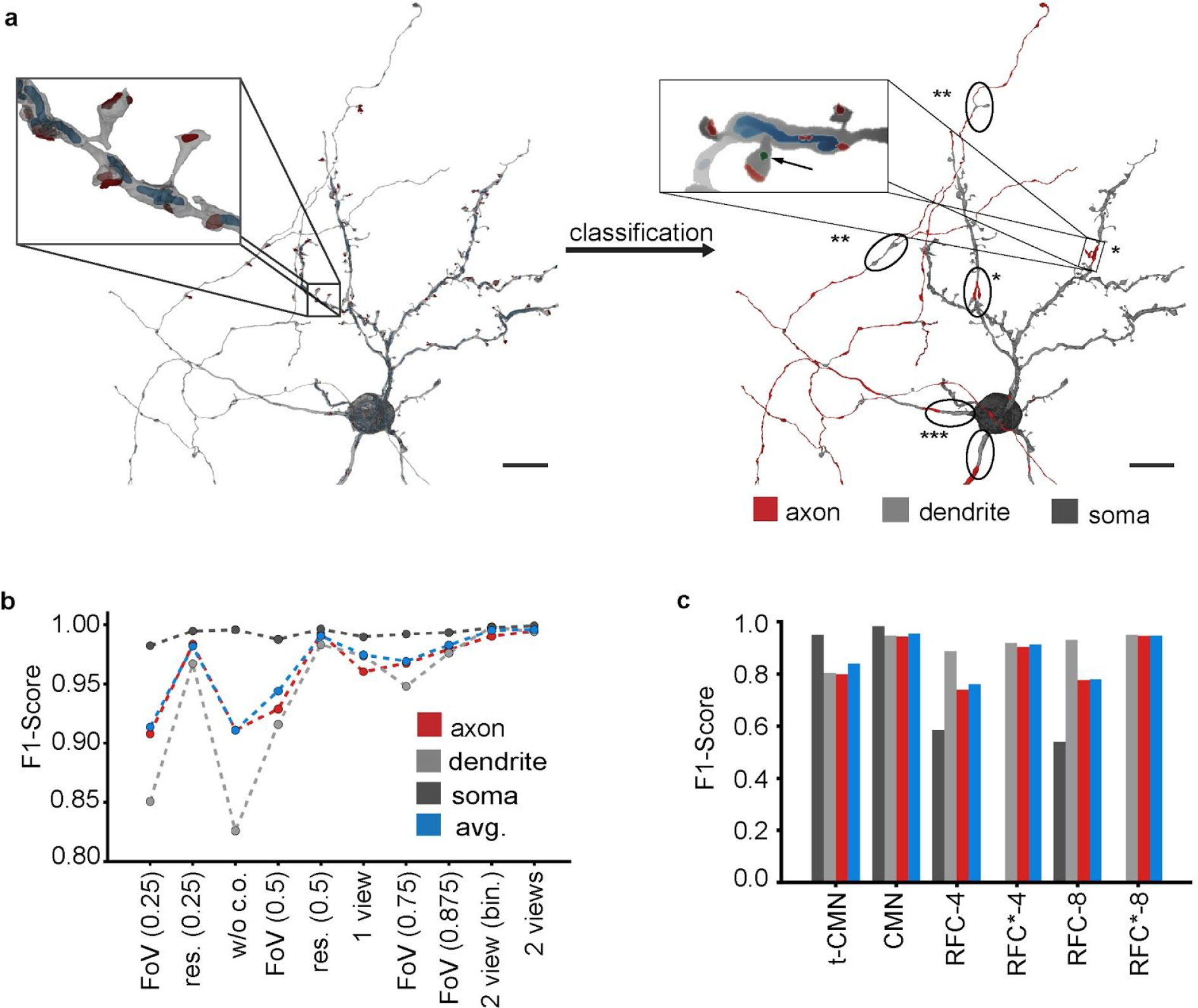
CMN-prediction of subcellular compartments. **a** The 3909 rendering locations of the reconstruction were predicted as axon, dendrite or soma. Local errors are indicated as follows: * indicate locations where vesicle clouds were falsely mapped to the cell (see inlay); ** indicate branch points in axons where synaptic junctions occurred without vesicle clouds; *** indicate false dendrite predictions at neurite outgrowths. **b** Performance on the validation set (axon: 1078; dendrite: 1823; soma: 6139 multi-views) for different inputs sorted by the number of input pixels. Left to right: FoV reduction by image cropping (3/8 on each side); resolution reduction by 4x downsampling; input views without cell object channels; FoV reduction by cropping (1/4); resolution reduction by 2x downsampling; only the view perpendicular to the 1st and 2nd p.c. was used; FoV reduction by cropping (1/8); FoV reduction by cropping (1/16); input views were binarized; both views were used at full resolution (256 × 128 px). **c** Comparison of skeleton and multi-view-based classifications measured on a skeleton node test set (color-code as in **b**) of 28 SSVs (N_axon_: 29,243; N_dendrite_: 38,170; N_soma_: 32,702). * indicates that the RFC model was trained on binary label data (axon vs. dendrite) only. The CMN model was the 2-view model evaluated in **b**; the t-CMN had N_r_=9, d_z_=10 and k=5. Scale bars are 10 μm.

### High-resolution semantic segmentation of neurite surfaces

The so far described classification of neurites is restricted in its spatial resolution to the minimum size of at least one view rendering (8×4×4 μm). This property makes the approach suitable for the identification of larger neuronal compartments (axons, dendrites and somata), which even benefits from large spatial context (Fig. 5b), but does not allow semantic segmentation of the morphology of a cell at sub-micron resolution. Higher resolution classification requires a dense analysis of the rendered views. Similar to the approach taken by Boulch et al.^32^, we solved the resulting image-to-surface mapping problem by rendering the cell views with spatially subdivided unique colors^33^, thereby creating an efficient reversible mapping between the 2D view space and the 3D surface (Fig. 6a-c). We then trained a VGG13 and FCN ^34,35^ based model on the identification of dendritic spines (Fig. 6d,e; the training set contained five MSN reconstructions; 12.59 mm; 1.01 GV; 1652 μm^3^), a classification problem that was previously solved on manually traced skeletons, which made the automated identification of spines easy ^11,36^. Automatically generated skeletons or surface meshes do not have the advantage that many skeleton endings are dendritic spines, which is likely a result of the implicit knowledge of human annotators. Instead of evaluating the classification performance on sampled locations, as done before by us^11^ (vertex-based evaluation in suppl. Text S1), we tested the performance on a test set of 182 manually annotated synaptic contacts, that were morphologically classified as a spinous synapse (N=88) or dendritic shaft synapse (N=94), F1-score 0.978 (prec. 0.978, recall 0.978, F1-score spine head only 0.977; 2 views per location and k=20). Interestingly, the classification performance did not improve further with more views (F1-score for k=20 and 6 views: 0.978), likely because the PCA view alignment along the dendritic process already optimized the coverage. Note that the coverage saturates below 1.0, due to non-surface triangles (see Fig. 6c).

**Fig. 6.**
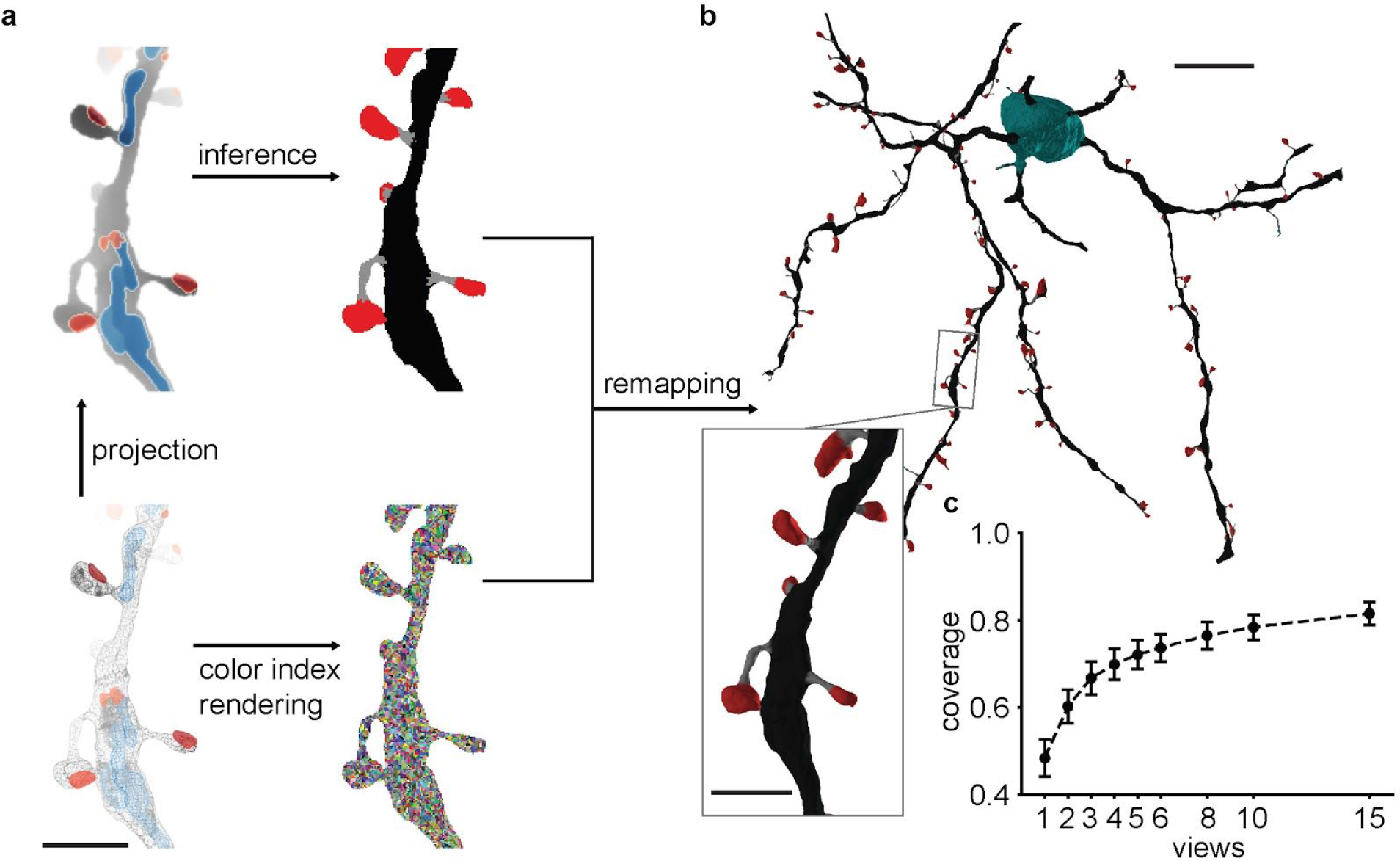
Sub-micron resolution semantic segmentation of cellular surfaces. **a** Bottom left: Wire-frame rendering of an FFN reconstructed mesh with mapped organelles. Top left: Depth map projection with organelle channels. Bottom right: Reversible 2D-to-3D mapping based on unique color rendering. Top right: Dense prediction of semantic pixel labels (here spine head (red), neck (grey), dendritic shaft (black)) on rendered view. **b** Pixel labels are mapped back onto mesh faces through the unique colors, enabling a high-resolution 3D surface classification. **c** Mean ratio of triangle faces covered by all reconstruction multi-views depending on the number of views rendered per location (error bars: s.d.). Scale bars in **a** are 2 μm, 10 μm in **b** and 2 μm in the inset.

## Discussion

We demonstrated that CMNs are a versatile tool to analyze reconstruction fragments or entire cell reconstructions. Our approach adapts to the neurons’ sparse coverage of the 3D volume by using distributed 2D multi-view projections instead of dense 3D models that end up processing many empty voxels^5,14,18,20^. While mainly developed for the analysis of neuron reconstructions, our proposed view-sampling method seems generally beneficial for the analysis of 3D objects that span large distances but still require high-resolution representations, as demonstrated by SnapNet which was developed independently for the semantic labeling of point clouds^32^.

By combining the concept of CMNs with unsupervised triplet loss training^22^, we created Neuron2Vec embeddings, that could serve as the basis for an unbiased morphological comparison of cells and cell types without requiring hand-designed features, or allow neuron-database queries using example neurites^16^. Additionally, the embedding can be used for visualization purposes e.g. through coloring (Fig. 2), making morphologically different regions salient. The triplet-loss embedding could be compared to alternative unsupervised training paradigms (e.g. autoencoders ^37,38^ or generative query networks^39^), to evaluate whether the excellent supervised training results can be reached. CMNs showed improved performance in cell compartment classification in comparison to the previously used hand-designed features^11^ and are likely more generalizable, since the morphology does not need to be parametrized first.

We used the classification results of the glia CMN for top-down neurite segmentation, where we evaluated so far only the splitting of falsely merged cells. However, the other identified cell types and neuronal compartments could be used in a similar way, or used as input to a graph cut segmentation algorithm, as proposed by Krasowski et al.^17^. Another application could be to directly evaluate the shape-plausibility of neurite fragments and detection of errors ^5,14^, or to estimate the probability that separate SVs should be combined^40^. Especially applications requiring high-resolution segmentation (e.g. the localization of reconstruction errors) should benefit from the cell surface analysis that we used here for the classification of postsynaptic dendritic morphology with excellent performance (F1-score 0.98). PointNets^21^ or PointCNNs^41^, which can operate directly on mesh vertex data, might be an alternative, but their effectiveness for neuronal morphology classification remains to be demonstrated and compared to the projection-based CMNs.

## Methods

### EM data and used segmentation

The analyzed EM data set (Area X, adult male zebra finch, >120 days post hatching) was acquired by JK through serial block-face scanning electron microscopy and had an extent of 96 × 98 × 114 μm^3^ with an xyz-resolution of 9 × 9 × 20 nm^3^. It contains about 664 GV (10664 × 10914 × 5701 voxel). Meshes and skeletons were based on a FFN segmentation by MJ and VJ^8^, including the over segmentation (all SVs) and the post-agglomeration SV-graph (defines SSVs).

### Local scene rendering

We used SV triangle meshes to efficiently render depth-maps with PyOpenGL (http://pyopengl.sourceforge.net/) and OSMesa (required for off screen rendering). For rendering, the modelview matrix was rotated such that the first axis of the view was parallel to the main principal component of the object and clipped to an extent of 8 × 4 × 4 μm^3^. The rendering had a throughput of about 3 multi-views/s (see Tab. S1) on a single CPU core, making it scalable in a cloud - or high-performance computing environment. We explicitly chose software rendering, since the view generation on the GPU was overall less economical, likely due to the overhead of main memory to GPU memory data transfers and higher availability of CPU cores.

PCA was applied to a subset (0.125) of the vertices within the clipping box which yielded the axes with highest variance (*x*, *y* and *z*; decreasing variance) in good approximation. In this system, *xy* and *xz* represent the planes with the highest spatial variance, allowing alignment with the elongated neurite structure. By rotating around the *x*-axis, orthographic depth-map projections of size 256 × 128 pixels were rendered and stored as unsigned 8-bit integer. For every SV, rendering locations were obtained by calculating its vertex density ρ in a grid of voxels with size 2 × 2 × 2 μm^3^. In order to not oversample dense regions, the center coordinate of every voxel with ρ > 0 was used to calculate the mean of the vertices within a radius of 1 μm. The resulting set of coordinates, effectively a downsampled point cloud, was stored as the SVs rendering locations.

### Rendering cell organelles

Three additional channels were generated next to the 2D depth-map views which contained the rendering of mapped cell organelles. Only cell organelles (mitochondrion: MI; synaptic junction: SJ; vesicle cloud: VC) with a relative overlap with SSVs above or within a certain threshold range (MI: > 0.5, VC: > 0.5, SJ: 0.2 − 0.8) were kept. Only objects with a minimum size (number of voxels) were taken into account when calculating the overlap (SJ: 498; VC: 1584; MI: 2786).

Meshes for the associated objects were extracted from a Gaussian-smoothed (σ = 1), distance-transformed (https://ukoethe.github.io/vigra/) binary 3D mask with marching cubes (contour value of 0; scikit-image http://scikit-image.org/).

The cell object meshes were rendered from the same perspective and resolution (256 × 128 px) as the corresponding SV views. The location’s fingerprint finally consisted of the rotated views, each with four channels.

### Automatic skeletonization of cell reconstructions

The skeletons of SV (provided by MJ and VJ and created using the TEASAR^42^ algorithm) belonging to an SSV were combined iteratively. Edges between the spatially closest pair of nodes of all connected components were repeatedly added until only a single connected component remained.

We decreased the average edge length in the skeleton representations to approx. 150 nm by removing skeleton nodes (ignoring branch and end nodes). Nodes of degree 2 were removed and replaced by a single edge if the summed length λ of the adjacent edges was below a threshold (*λ*≤50 nm) or if the dot product of their edges was higher than 0.8 in combination with *λ* < 500 nm.

At every node, the cell radius was estimated by the median of the distance to the ten nearest vertices of the mesh. Total path lengths were calculated as the sum of all edges.

### Multi-view models for type classification

Multi-views were sampled from the joint set of SV meshes of anentire SSV, and renderings generated at locations as described above.

To overcome RAM limitations, large SSVs (>10^4^ SVs) were processed as subgraphs, defined by a breadth-first-search (extending 40 nodes) on the SV graph and starting at each SV.

Multi-views were generated from the joint meshes of the 40 SVs at the sampled locations of the source SV.

The multi-view CNNs used seven convolutional layers (number of filters, filter size, max-pooling size), each followed by a max-pooling layer, three fully connected layers and a soft-max layer:

- conv. L1: (13, (1, 5, 5), (1, 2, 2))
- conv. L2: (17, (1, 5, 5), (1, 2, 2))
- conv. L3: (21, (1, 4, 4), (1, 2, 2))
- conv. L4: (25, (1, 4, 4), (1, 2, 2))
- conv. L5: (29, (1, 2, 2), (1, 2, 2))
- conv. L6: (30, (1, 1, 1), (1, 2, 2))
- conv. L7: (31, (1, 1, 1), (1, 1, 1))
- f.c. L1: 50 neurons
- f.c. L2: 30 neurons
- f.c. L3: 3 neurons
- softmax-layer

The input to a model contained multi-views with either one or four channels (c) and with shape ((b, 20), (f, c), (z, 2), (x, 256), (y, 128)), with batch size b, initial filter or channels f and spatial axes z, x and y. Note the auxiliary z-axis which was introduced to share filters for all views (z-filter size of 1). The models were trained using backpropagation with mini-batches (samples were drawn uniformly). During training, the ordering of the two views was inverted with probability *p* = 0.5.

If not stated otherwise, hyper-parameters were chosen to be:

- batch size: 20
- dropout rate ^43^: 0.1
- activation function: ReLu
- Adam optimizer ^44^
- learning rate: 1 · 10^−4^
- momentum β _1,2_: 0.9, 0.99
- weight decay: 5 · 10^−4^

A model with two output classes and one input channel had 35,770 trainable parameters and a computational cost of 2.4 Giga Ops. 350,000 training iterations (with batch size of 20) on a GeForce GTX 980Ti took 20.1h, which was about 0.0103s per sample. Inference of 2,182 samples took 16.33s, approx. 0.0075s per sample (see Table S1).

### Neuron2vec embedding

The architecture of the CMN encoder was used to learn a projection from the single view space ℝ^256·128·4^ to a lower dimensional latent space (embedding) ℝ^*d*^ (d dimensions) based on the triplet loss described in ^23^. Its architecture was identical to the one described in “Multi-view models for type classification”, whereas max pooling was removed from conv. L2 and L6, dropout was restricted to conv. L3-L5, the f.c. L2 had 50 hidden units and the softmax layer was removed. The objective function was to keep the distance of the reference *x*_*ref*_ to the similar input *x*_+_ below the distance of *x*_*ref*_ to the dissimilar input *x*_−_. For the two similar views *x*_*ref*_ and *x*_+_ we used the two views of the same rendering location (rotated by φ=50°), while the dissimilar view *x*_−_ was sampled randomly from a different SSV. The clipping volume was set to 8 × 4 × 8 μm^3^. In order to take strong similarities of adjacent rendering locations into account, we modified the view-sampling during training and allowed to sample similar views also from close-by rendering locations instead of only using the rotated view at the same rendering location (Fig. S2).

The loss was defined as *L*_α>0_ = α + λ_2_ and *L*_α≤0_ = λ_2_, with α = *r*_+_ − *r*_−_ + λ_1_, *r*_+_ = | *x̃*_*ref*_ − *x̃*_+_|_2_ *r*_−_ = |*x̃_ref_* − *x̃*_−_|_2_ and λ_1_ = 0.2 being a parameter to control the margin between data points. *x̃* ⊂ ℝ^*d*^ represents the latent space of the triplet net. The second regularization term λ_2_ was the mean norm of the reference, similar and dissimilar view and acted as a counterpart to λ 1 by restricting the latent vectors to be small: λ_2_ = 1/3 (|*x̃_ref_* |_2_ + |*x̃*_+_|_2_ + |*x̃*_−_|_2_). Our implementation was inspired by the one from A. Veit (https://github.com/andreasveit/triplet-network-pytorch). The PCA was performed on the triplet network latent vectors of the 372 cell or cell fragment reconstructions which were also used for querying the view-triplets during training. By taking only the first three principal components, every multi-view location was assigned an RGB value. The mesh was colored according to the nearest view color of every vertex.

### Glia classification

For the glia classification model, only the depth maps of the SV were given as input. The training set contained 88,022 multi-views (N_neuron_: 69,068; N_glia_: 18,954) and the validation set 9,695 (N_neuron_: 7588; N_glia_: 2107). Neuron views from the subcellular compartment ground truth (see next section) were extended by two additional axon reconstructions and used as samples for the negative class (31.29 mm 3.08 GV; 4989 μm^3^). Glia views were generated from 118 manually annotated glia SV (path length: 337.45 mm; 13.40 GV; 21706 μm^3^). The performance was calculated based on multi-views and measured as F1-score. The classification threshold θ was set to the optimal F1-score on the validation set. The best model was retrained on training and validation set and applied to the whole data set. To remove background structures not connected to the central object of interest, connected component analysis was performed on the 2D multi-view images, followed by masking of the unconnected pixels.

The SVs were then classified by calculating the mean of all its multi-view predictions and thresholding with θ. In addition, at least 70% of all multi-view predictions of a SV had to be glia for the assignment of this label.

Classification performance was measured by manually annotating 169 SV (N_glial_: 85; N_neuron_: 84; N_BBD<8μm_: 57; N_BBD ≥ 8μm_: 112). These SVs were sampled from 20,000 randomly drawn SSVs (training and validation samples were excluded), weighted by the number of views per SV. Only SSVs within the segmentation data set bounding box [470, 730, 30] to [10200, 10200, 5670] were taken into account.

### Top-down glia splitting

In order to split glia fragments from neurons, a CC analysis was applied to the glia and neuron SV-graphs identified by the SV glia predictions of every SSV. The glia and neuron CC size was estimated by calculating their BBD and glia CCs with a BBD ≥ 8.0 μm were removed from the SV-graph first. The remaining, small glia CCs (BBD<8.0 μm) were assigned the neuron class and the BBD was re-evaluated. Neuron CCs with a BBD below 8 μm were removed and added to the glia graph. The purpose of this was to bridge small false glia/neuron predictions and thereby avoid false splits.

Splitting performance was evaluated on twelve randomly drawn SSVs with at least one split introduced during the splitting procedure. Inspected SVs were sorted by volume and the average inspected volume coverage was 0.905 (proportion of inspected SVs weighted by their volume).

### Neuron type classification

The 2-views were re-used to construct the N-views by the following procedure: The collection of all M 2-views of an SSV was split into 2*M*/*N* random sets (drawn without replacement) each of size *N*. If 2*M*< *N*, the set was filled by randomly drawing from the existing views. The model architecture was identical to the model used for glia classification, except for a reduced batch size, a dropout rate of 0.08 and a learning rate schedule defined as exponential decay, with decay rate of 0.98 per 1000 steps. The input shape was (1, 4, N, 256, 128).

We used 402 manually traced (skeletonized) cells to identify their corresponding SSV which were split into training (N_train_: 301; N_EA_: 177, N_MSN_: 114, N_GP_: 6, N_INT_: 6) and test set (N_test_: 101; N_EA_: 60, N_MSN_: 39, N_GP_: 3, N_INT_: 2) with the following labels, that correspond to the broad biological classes found in this data set (excitatory axons (EA), medium spiny neurons (MSN), pallidal like neurons (GP), interneurons (INT)).

During batch creation while training, the N-views were generated by randomly drawing from the corresponding SSV views. Every batch contained an equal number of SSVs for each class. The classification was performed using arg-max on the output of the softmax layer and the majority vote of the corresponding N-view classifications was used for SSV classification. Performance was evaluated on N-views and on a SSV level after majority vote. The latter was measured as unweighted F1-score of the classes EA and MSN due to the little support for INT and GP (EA: 60, MSN: 39, GP: 3, INT: 2).

### Subcellular compartment classification

The cellular compartments of 33 neurites were manually annotated and axon, dendrite and soma views generated, which were split into a training set (N_train_: 80,370 views; N_dendrite_: 10,004; N_axon_: 41,424; N_soma_: 28,942) and a validation set (N_validation_: 9,040; N_dendrite_: 1,078; N_axon_: 1,823; N_soma_: 6139). During training we applied class weights for loss computation to address imbalances in their frequency (dendrite: 2, axon: 1, soma: 1). Performance was measured with the F1-score of the multi-view classification using argmax on the softmax output. The best model was again retrained on the whole ground truth data for the data set prediction.

Classification of arbitrary locations within neurons was performed by assigning the unclassified location the label of the closest classified location (Voronoi partitioning with Euclidean distance).

#### Comparison with RFC

All SSV skeleton nodes were manually labeled (N_axon_: 29,243; N_dendrite_: 38,170; N_soma_: 32,702) by a human expert using KNOSSOS. In order to enable a direct comparison between the two models, the skeleton-node locations were used for the extraction of the hand-designed features. CMN and kNN predictions were mapped to the skeleton nodes using the nearest neighbors on the multi-view locations. As in ^11^, hand-designed features were computed for every skeleton node (context of 4,000 nm and 8,000 nm maximal traversed path length from the source node). Only properties of nodes visited during the traversal were taken into account for the source node statistics.

A total of 23 features were extracted from the collected properties at each node: Mean and standard deviation (s.d.) and histogram (10 bins) of the encountered node diameters, mean of node degrees, node density within a box with edge length of 2-times the context-range (either 4μm or 8μm) number of cell organelles and mean and s.d. of their size for mitochondria, synaptic junctions, vesicle clouds. The RFC was trained on the same training data as the CMN to classify each node as axon, dendrite or soma using argmax on the resulting class probability.

A majority vote was carried out on the set of node/multi-view predictions collected within a 12.5 μm traversal depth for every skeleton node for the RFC and CMN/kNN, respectively.

### High-resolution semantic segmentation of surfaces

The training data was generated by rendering multi-views (5 different perspectives) from the rendering locations of 5 reconstructions (training: 24,248 views; validation: 6,062 views) with label-dependent vertex colors. Skeleton nodes were manually annotated as either neck, head, shaft or soma/axon, which were then mapped to the mesh vertices with Voronoi-partitioning (Euclidean distance). To smooth label boundaries, each vertex was assigned the majority label of 40 vertices found by a BFS on the vertex graph of the reconstruction. Graph edges were added between vertices with a distance of up to 120 nm. Only rendering locations with annotated skeleton nodes within 2 μm were taken into account.

We used the FCN-VGG13 architecture^35^ (adopted from https://github.com/pochih/FCN-pytorch) to perform pixel-wise multi-class (neck, head, shaft, background and axon/soma) classification on single views with four channels (cell, mitochondria, synaptic junctions and vesicle cloud shapes). It was trained using backpropagation with mini-batches (images were flipped in x/y with probability *p* = 0.5), Adam optimizer (β_1,2_: 0.9, 0.999; weight decay: 5 · 10^−4^), initial learning rate of 4 · 10^−3^ (exponential decay with 0.99) and Lovász-Softmax loss^45^.

To assign pixel labels back to the mesh vertices an additional view of the faces was rendered using color buffering with a unique ID per face. This allowed to perform a majority vote of all collected labels corresponding to a single vertex as classification. Subsequently, we applied a kNN classification to propagate predicted labels to vertices which were not covered by the rendered color map.

The synapse test set was generated as a random set of head and shaft synapses collected from 4 different reconstructions. The dendritic tree for the per-vertex evaluation was manually annotated on skeleton node level which were then propagated to the mesh vertices as described above for the GT generation.

### Reconstructing glial cells

SV graph edges were added between the sample locations of the collection of all splitted glia SV by identifying the k-nearest neighbors (k=15, maximum distance: 10 μm; weighted by euclidean distance).

27 somata of putative astrocytes were identified in the data set and every glia SV was assigned to its closest soma (shortest path using Dijkstra’s algorithm).

### Bloodvessel prediction

The input data (zxy ordering) for the bloodvessel CNN was downsampled by a factor of 8 in all dimensions. A cube of size (256, 437, 287) was densely labeled using KNOSSOS to obtain training data. The used network had the following architecture:

- conv. L1: (24, (1, 6, 6), (1, 2, 2))
- conv. L2: (27, (1, 5, 5), (1, 2, 2))
- conv. L3: (30, (1, 5, 5), (1, 1, 1))
- conv. L4: (33, (1, 4, 4), (2, 1, 1))
- conv. L5: (36, (3, 4, 4), (1, 1, 1))
- conv. L6: (39, (3, 4, 4), (1, 1, 1))
- conv. L7: (42, (2, 4, 4), (1, 1, 1))
- conv. L8: (45, (1, 4, 4), (1, 1, 1))
- conv. L9: (48, (1, 4, 4), (1, 1, 1))
- conv. L10: (48, (1, 1, 1), (1, 1, 1))
- conv. L11: (2, (1, 1, 1), (1, 1, 1))
- softmax-layer

The dense predictions of the dataset were thresholded at 0.98. Meshes were created as described above (“Rendering cell organelles”).

### Computing infrastructure

The used parallel computing environment consisted of 18 nodes, each equipped with 20 cores (Intel(R) Xeon(R) CPU E5-2660 v3 @ 2.60GHz), 2 GeForce GTX 980Ti and 256 GB of RAM. Compute jobs were managed using openmp in combination with SGE QSUB.

## Code availability

The used network architectures, classes for handling the inference, processing and storage of the segmentation data can be found in the SyConn GitHub repository (https://github.com/StructuralNeurobiologyLab/SyConn/).

## Acknowledgements

We would like to thank W. Denk for enabling this work in his department and many helpful discussions, M. Fee and M. Kormacheva for helpful comments, the KNOSSOS team for support, R. Saxena and M. Shumliakivska for code contributions, M. Killinger and M. Drawitsch for help with the ELEKTRONN library and J. Rogowska and M. Shumliakivska for proofreading tracings and ground truth generation.

## Author contributions statement

PS performed and designed experiments and wrote the manuscript.

MJ and VJ contributed the data set FFN segmentation and wrote the manuscript. SD contributed code for the experiments.

JK designed experiments and wrote the manuscript.

